# Regulation of cell size and Wee1 by elevated levels of Cdr2

**DOI:** 10.1101/2022.07.27.501696

**Authors:** Rachel A. Berg, James B. Moseley

**Author notes:** Correspondence to James B. Moseley.

## Abstract

Many cell cycle regulatory proteins catalyze cell cycle progression in a concentration-dependent manner. In fission yeast *S. pombe*, the protein kinase Cdr2 promotes mitotic entry by organizing cortical oligomeric nodes that lead to inhibition of Wee1, which itself inhibits Cdk1. _*cdr2*_*Δ* cells lack nodes and divide at increased size due to overactive Wee1, but it has not been known how increased Cdr2 levels might impact Wee1 and cell size. Using a _Tet_racycline-inducible expression system, we found that a 6X increase in Cdr2 expression caused hyperphosphorylation of Wee1 and reduction in cell size. This overexpressed Cdr2 formed clusters that sequestered Wee1 adjacent to the nuclear envelope. Cdr2 mutants that disrupt either kinase activity or clustering ability failed to sequester Wee1 and to reduce cell size. We propose that Cdr2 acts as a dosage-dependent regulator of cell size by sequestering its substrate Wee1 away from Cdk1 in the nucleus. This mechanism has implications for other clustered kinases, which may act similarly by sequestering substrates.

## Introduction

The core elements of the eukaryotic cell cycle include a regulatory network that promotes switch-like entry into mitosis at the G2/M transition. Activated cyclin-dependent kinase Cdk1 bound to its cyclin subunit phosphorylates diverse substrates to drive mitotic entry (Morgan, 1997; Wieser and Pines, 2015). In G2, the Cdk1-cyclin complex is kept inactive by Wee1 kinase, which inhibits Cdk1 by phosphorylating a conserved tyrosine residue (Coleman and Dunphy, 1994; Kellogg, 2003). At the G2/M transition, Cdk1 inhibition is reversed by the phosphatase Cdc25, which removes the inhibitory tyrosine phosphorylation from Cdk1 (Millar and Russell, 1992; Coleman and Dunphy, 1994). Activated Cdk1-cyclin then inhibits Wee1 and activates Cdc25, resulting in dual feedback loops for switch-like mitotic entry (Wieser and Pines, 2015). In the fission yeast *Schizosaccharomyces pombe*, a long-standing model organism for cell cycle research, regulation of these conserved cell cycle proteins establishes cell size at division. More specifically, fission yeast cells enter mitosis at a reproducible size due to size-dependent activation of Cdk1 regulated in part by Wee1 and Cdc25 (Rupes, 2002).

Many of these proteins act as dosage-dependent regulators of fission yeast cell size at division, consistent with their known activities and mechanisms. For example, loss-of-function mutations in *wee1+* cause cells to divide at a small size due to over-active Cdk1, while *wee1+* over-expression increases cell size at division (Russell and Nurse, 1987a). Conversely, mutations in *cdc25+* increase cell size, while *cdc25+* overexpression reduces cell size (Russell and Nurse, 1986). Wee1 activity in cells is regulated in a size-dependent manner by the protein kinases Cdr1 and Cdr2 (Allard *et al*., 2018), which are conserved SAD family kinases. Cdr1 (also called Nim1) directly phosphorylates Wee1 to inhibit its kinase activity (Coleman *et al*., 1993; Parker *et al*., 1993; Wu and Russell, 1993; Opalko *et al*., 2019). Cdr1 acts in a dosage-dependent manner: loss-of-function causes elongated cells due to over-active Wee1, while *cdr1*+ overexpression reduces cell size at division (Russell and Nurse, 1987b). These results are consistent with Cdc25, Wee1, and Cdr1 acting as concentration-dependent catalytic regulators of their substrates.

Cdr2 also promotes Wee1 inhibition in cells but does not appear to inhibit Wee1 kinase activity directly (Kanoh and Russell, 1998). Rather, Cdr2 forms oligomeric “nodes” at the plasma membrane and recruits both Cdr1 and Wee1 to these structures (Morrell *et al*., 2004; Martin and Berthelot-Grosjean, 2009; Moseley *et al*., 2009; Allard *et al*., 2018). Consistent with this mechanism, loss-of-function mutations in _*cdr2*_*+* increase cell size (Young and Fantes, 1987; Breeding *et al*., 1998; Kanoh and Russell, 1998). However, _*cdr2*_*+* overexpression has led to mixed results, leaving it unclear how this protein acts on cell size in a dosage-dependent manner. Strong *cdr2+* overexpression with the *P3*_*nmt1*_ promoter is lethal, and cells exhibit pleiotropic defects including multi-septation and branching (Breeding *et al*., 1998). *P3*_*nmt1*_*-cdr2+* overexpression causes a shift in the migration of Wee1 by Western blot (Opalko *et al*., 2019), but the lethality of this strong overexpression system prevented further mechanistic and functional studies. Lower levels of *cdr2+* overexpression with the weakened *P81*_*nmt1*_ reduce cell size (Bhatia *et al*., 2014; Pan *et al*., 2014), but the underlying mechanism has not been studied.

Here, we increased _*cdr2*_*+* expression with a _Tet_racycline-regulated promoter recently adapted for controlled expression in *S. pombe* (Patterson *et al*., 2019). We discovered that increased expression of _*cdr2*_*+* induces Wee1 hyperphosphorylation and sequestration at cytoplasmic clusters, leading to premature mitotic entry and reduced cell size at division. These findings establish Cdr2 as a dosage-dependent regulator of cell size through a localization-based mechanism.

## Results and Discussion

We used the _Tet_racycline-induced expression system to control levels of Cdr2. In the presence of anhydrotetracycline (Tet), *P*_*Tet*_*-GST-cdr2* (hereafter *P*_*Tet*_*-cdr2*) integrated at the *leu1+* locus slowed the migration of Wee1-FLAG by SDS-PAGE, but catalytically inactive mutants *P*_*Tet*_*-cdr2(E177A)* and *P*_*Tet*_*-cdr2(T166A)* did not affect Wee1-FLAG (Figure 1A and S1A). We confirmed that *P*_*Tet*_*-cdr2* induced a similar shift for untagged Wee1 (Figure S1B). This band shift was due to hyperphosphorylation because it was reversed by treatment with λ-phosphatase (Figure 1B). To determine the level of overexpression responsible for this effect, we generated a *P*_*Tet*_*-cdr2-5FLAG* construct for comparison with _*cdr2*_*-5FLAG* expressed by the endogenous promoter at the endogenous chromosomal locus. By quantitative Western blot of whole cell lysates, the level of *P*_*Tet*_*-cdr2-5FLAG* was 6 times higher than endogenously expressed *cdr2-5FLAG* (Figure 1C). We considered that hyperphosphorylation of Wee1 might involve the presence of endogenously expressed Cdr1 and Cdr2 in this system, so we repeated this experiment in *cdr1Δ* _*cdr2*_*Δ* cells. Interestingly, *P*_*Tet*_*-cdr2* still induced hyperphosphorylation of Wee1 in *cdr1Δ cdr2Δ* cells (Figures 1D and S1C), showing that this modification is not mediated by Cdr1.

**Figure 1.**
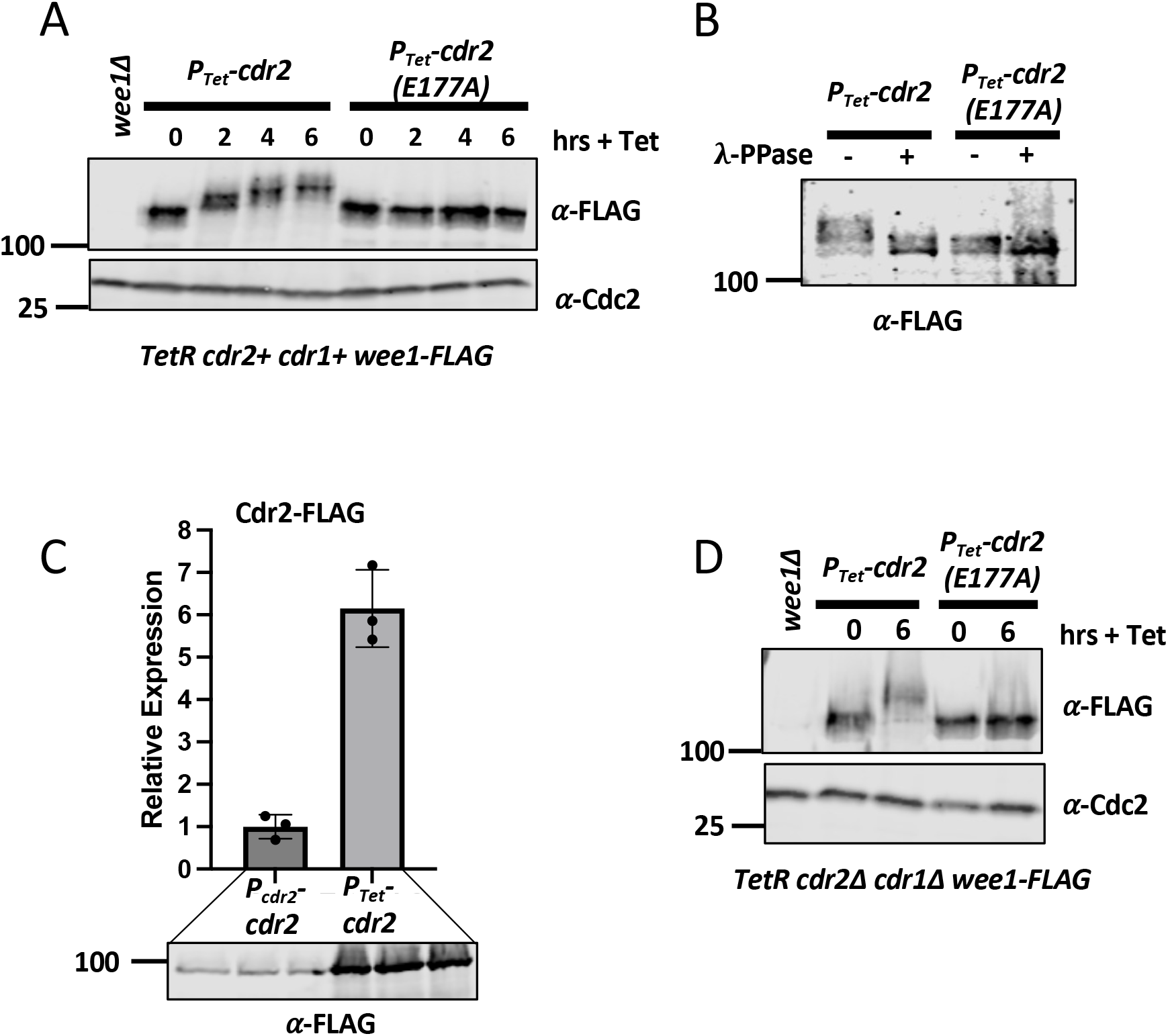
Tet-regulated Cdr2 overexpression induces hyper-phosphorylation of Wee1. (A) Whole-cell extracts were separated by SDS-PAGE and Western blotted to detect Wee1-FLAG after induction of wild type Cdr2 or kinase-dead Cdr2(E177A). (B) Gel shift induced by Cdr2 over-expression is reversed by А-phosphatase. (C) Western blot quantification of cellular Cdr2-FLAG protein overexpression. (D) Hyperphosphorylation of Wee1 does not require endogenous Cdr1 or Cdr2. Cdc2 was probed as a loading control.

Next, we tested the effects of *P*_*Tet*_*-cdr2* on cell size. Addition of Tet to *P*_*Tet*_*-cdr2* cells caused a marked and significant decrease in cell length at division (Figure 2A-B). In contrast, _Tet_-based overexpression of _*cdr2*_*(E177A)* increased the size of dividing cells, consistent with dominant-negative effects for this inactive mutant. Cdr2 controls cell size by recruiting Wee1 to cortical nodes where it is inhibited by Cdr1 (Allard *et al*., 2018; Opalko *et al*., 2019), so we considered that *P*_*Tet*_*-cdr2* effects on cell size might require Cdr1. However, induction of *P*_*Tet*_*-cdr2* in *cdr1Δ cdr2Δ* cells still reduced cell size, whereas *P*_*Tet*_*-cdr2(E177A)* had no effect (Figure 2C-D). These results indicate that *P*_*Tet*_*-cdr2* induces Wee1 hyperphosphorylation and reduces cell size by a mechanism that is independent of Cdr1.

**Figure 2.**
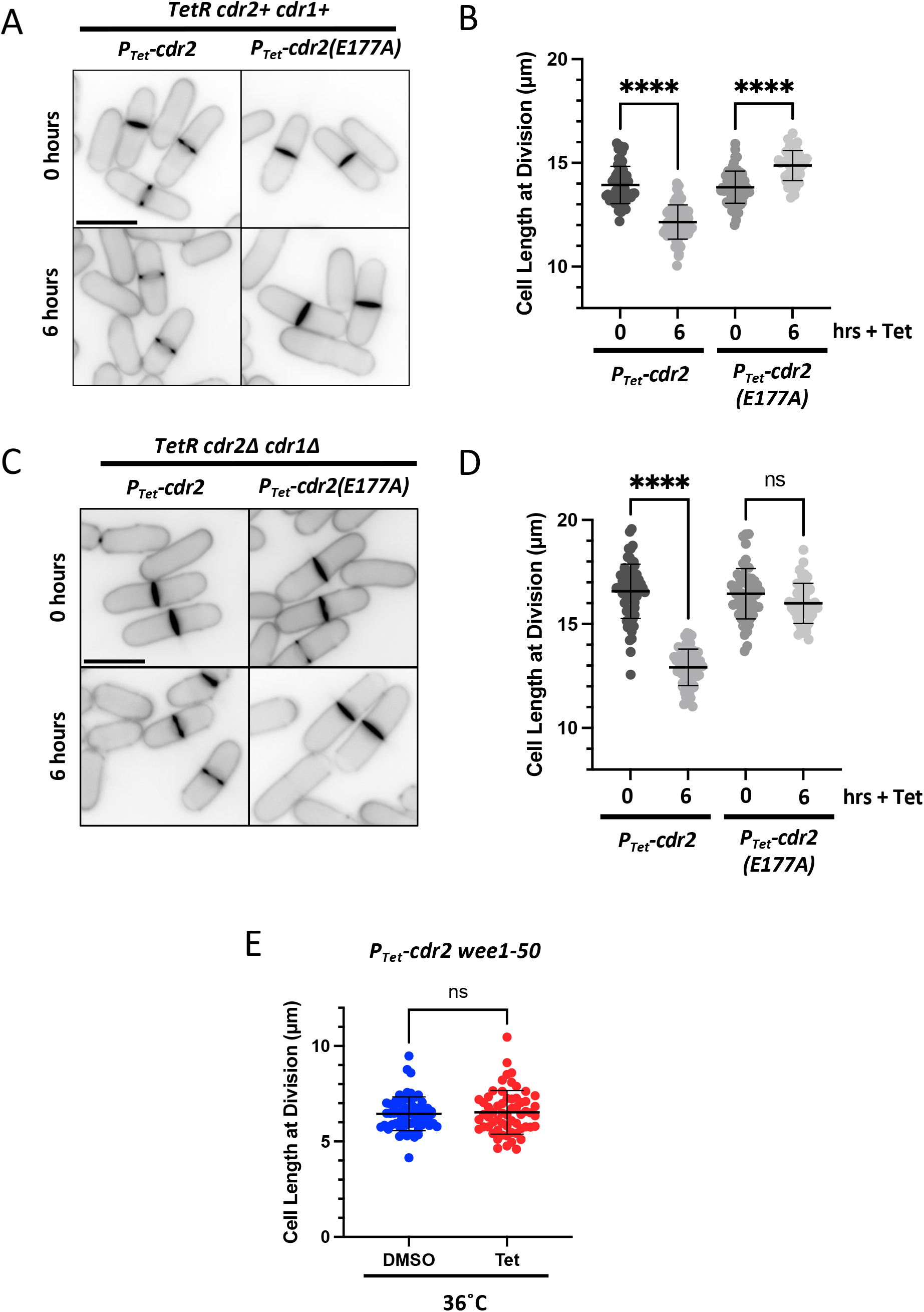
*P*_*Tet*_*-cdr2* reduces cell size at division. (A) Blankophor-stained images of cells before and after _Tet_ treatment. Scale bar, 10 µm. (B) Cell length at division for the indicated strains and treatments. n ≥ 50 cells each. (C) Blankophor-stained images of *cdr1Δ* _*cdr2*_*Δ* cells before and after _Tet_ treatment. Scale bar, 10 µm. (D) Cell length at division for the indicated strains and treatments. n ≥ 50 cells each. (E) Cell length at division for *P*_*Tet*_*-cdr2 wee1-50* cells grown at the nonpermissive temperature of 36°C for 4 hours. ns, not significant (p > 0.05). ****, p < 0.0001 determined by ANOVA (panels B and D) or Welch’s t test (panel E).

We used genetic epistasis to investigate the underlying pathway. If *P*_*Tet*_*-cdr2* regulates cell size through Wee1, then it should have no effect in the absence of Wee1 activity. Consistent with this prediction, *P*_*Tet*_*-cdr2* did not reduce cell size in the temperature-sensitive *wee1-50* mutant grown at 36°C (Figure 2E). We conclude that increased levels of Cdr2 cause hyperphosphorylation of Wee1 leading to reduced cell size at division.

To determine the mechanism for *P*_*Tet*_*-cdr2* regulation of Wee1, we examined the localization of mEGFP-Cdr2 induced by *P*_*Tet*_. Previous studies have shown that Cdr2 exerts spatial control over the Wee1 regulatory pathway. In addition to cortical nodes along cell sides, we observed bright cytoplasmic puncta for both mEGFP-Cdr2 and mEGFP-Cdr2(E177A) expressed by the *P*_*Tet*_ promoter (Figure 3A). Interestingly, induction of *P*_*Tet*_*-cdr2* caused recruitment of Wee1-mNG to similar cytoplasmic clusters (Figure 3B). This redistribution of Wee1 required Cdr2 kinase activity because it was lost in the *P*_*Tet*_*-cdr2(E177A)* mutant (Figure 3B).

**Figure 3.**
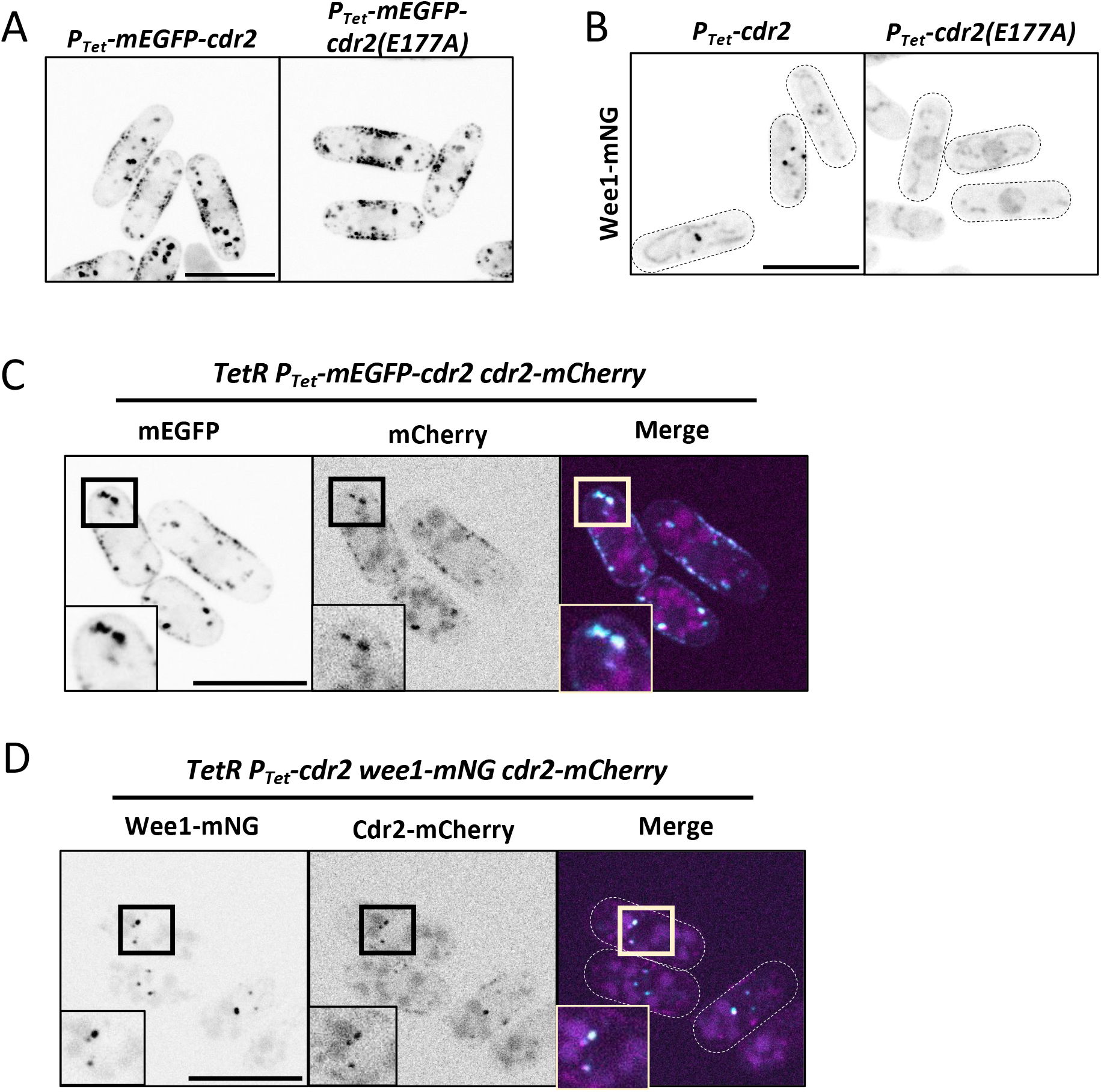
Cdr2 and Wee1 localize to cytoplasmic puncta in *P*_*Tet*_*-cdr2* induced conditions. (A) Images of the indicated strains after incubation with _Tet_ for 6 hours. Maximum intensity projections of 0.5 µm-spaced focal planes covering 2 µm in cell middle. (B) Wee1-mNG in the indicated strains after incubation with _Tet_ for 6 hours. Single focal plane. Cell boundaries are outlined. (C) Colocalization of P_Tet_-mEGFP-Cdr2 and P_cdr2_-Cdr2-mCherry after incubation with Tet for 6 hours. Maximum intensity projections of 0.5 µm-spaced focal planes covering 1 µm in cell middle. (D) Colocalization of P_cdr2_-Cdr2-mCherry and Wee1-mNG after Tet-induced expression of *P*_*Tet*_*-cdr2*. Maximum intensity projections of 0.5 µm-spaced focal planes covering 1 µm in cell middle. All scale bars, 10 µm. Boxed regions are zoomed in lower corner panels. Single channel images are shown with inverted LUT.

Next, we performed a series of colocalization experiments on these *P*_*Tet*_*-mEGFP-cdr2* cytoplasmic clusters. Upon induction, *P*_*Tet*_*-mEGFP-cdr2* recruited endogenously expressed Cdr2-mCherry to cytoplasmic clusters (Figure 3C), and this effect was independent of Cdr2 kinase activity (Figure S2A). Using endogenously expressed Cdr2-mCherry as a marker for clusters, we found that Wee1-mNG and Cdr2-mCherry colocalized in the same cytoplasmic clusters when expression of *P*_*Tet*_*-cdr2* was induced from the *leu1+* locus (Figure 3D). In contrast, the inactive mutant *P*_*Tet*_*-cdr2(E177A)* did not drive colocalization of Wee1-mNG and Cdr2(E177A)-mCherry at cytoplasmic clusters (Figure S2B).

Together, our results show that both active Cdr2 and inactive Cdr2(E177A) localize to cytoplasmic clusters upon expression with *P*_*Tet*_. However, only active Cdr2 clusters can recruit Wee1, induce Wee1 hyperphosphorylation, and reduce cell size at division. These results lead to a simple model in which Cdr2 clusters sequester Wee1 away from its nuclear target Cdk1, causing premature entry into mitosis.

Based on this working model, we sought additional information about these Cdr2 cytoplasmic clusters. Cdr2 recruits Cdr1 kinase and Arf6 GTPase to cortical nodes (Martin and Berthelot-Grosjean, 2009; Moseley *et al*., 2009; Opalko *et al*., 2022), but we did not observe strong recruitment of either Cdr1 or Arf6 to cytoplasmic clusters in *P*_*Tet*_*-cdr2* cells (Figure 4A-B). Thus, Cdr2 and Wee1 might be the primary components of these clusters. To probe the nature of Cdr2 interactions within clusters, we treated cells with 1,6-hexanediol, which disrupts weak, hydrophobic molecular interactions. Cdr2 cytoplasmic clusters disappeared after 10 minutes of treatment with 5% hexanediol (Figure 4C), indicating that these clusters are held together by weak interactions and are not irreversibly aggregated. Interestingly, endogenous Cdr2 nodes also were dispersed by the same 1,6-hexanediol treatment (Figure S2C). This result suggests that Cdr2 cytoplasmic clusters and endogenous Cdr2 nodes are held together by similar biophysical properties.

**Figure 4.**
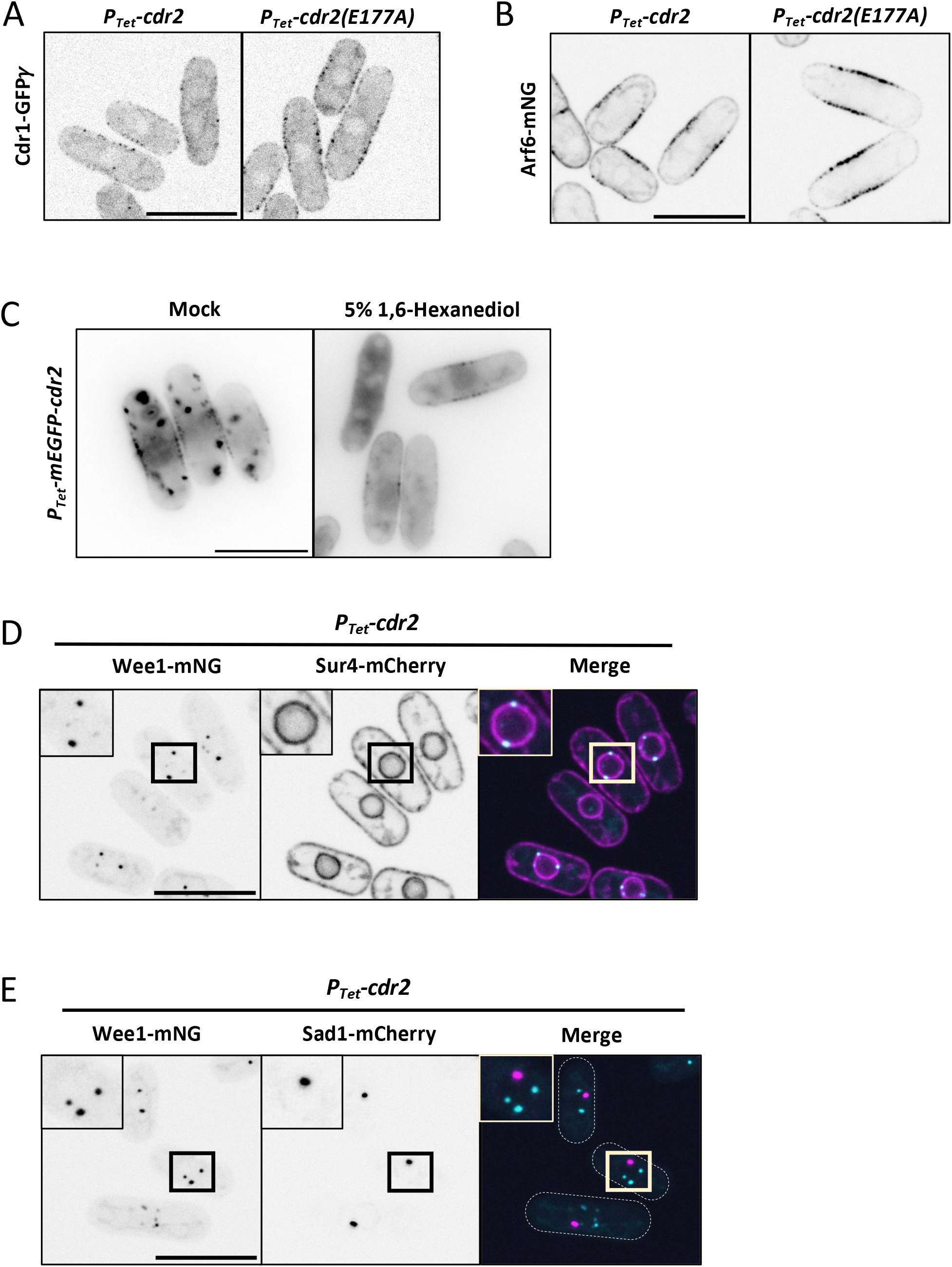
Characterization of Cdr2 clusters. (A) Localization of Cdr1-GFP*γ*in the indicated strains. (B) Arf6-mNG imaged after *P*_*Tet*_*-cdr2* and *P*_*Tet*_*-cdr2(E177A)* induction. (C) Induced *P*_*Tet*_*-mEGFP-cdr2* cells were treated with 5% 1,6-hexanediol. Middle focal plane images; scale bar is 10 µm. (D) Wee1-mNG and Sur4-mCherry after *P*_*Tet*_*-cdr2* induction. (E) Wee1-mNG and Sad1-mCherry after *P*_*Tet*_*-cdr2* induction. All scale bars are 10 µm. All images are maximum intensity projections from 0.5 µm-spaced focal planes spanning 1 µm in cell middle. Single channel images are shown with inverted LUT.

mEGFP-Cdr2 clusters can be found throughout the cytoplasm, but we noted that Wee1-mNG clusters in *P*_*Tet*_*-cdr2* cells are restricted to the cell middle and absent from cell ends. Using the ER marker Sur4-mCherry and the nuclear envelope marker Cut11-mCherry, we found that Wee1-mNG clusters in *P*_*Tet*_*-cdr2* cells are restricted to the nuclear periphery and/or nuclear ER (Figures 4D and S2D). We considered the possibility that these clusters might be the spindle pole body (SPB), which is embedded in the nuclear envelope, but Wee1-mNG clusters did not colocalize with the SPB marker Sad1-mCherry (Figure 4E; no colocalization in 20 of 22 cells examined). These results show that Cdr2 clusters can be found throughout the cytoplasm, but only clusters associated with the nucleus can capture and sequester Wee1.

Finally, we used the *P*_*Tet*_ system to define the domains of Cdr2 required for cluster formation and effects on both Wee1 and cell size. The Cdr2 N-terminus contains a kinase domain and a ubiquitin-associated (UBA) domain, which is thought to be necessary for kinase activity (Guzmán-Vendrell *et al*., 2015) (Figure 5A). The Cdr2 C-terminus contains a KA1 domain (kinase associated-1) and a basic patch, both of which promote binding to membranes (Rincon *et al*., 2014). Between the N-terminus and C-terminus is a linker domain that is predicted to be unstructured. Using *P*_*Tet*_ to drive expression, neither Cdr2(1-330) nor Cdr2(1-590) truncations reduced cell size or formed cytoplasmic clusters (Figure 5B-D). This result suggests that clustering mediated by the KA1 domain is required for reduction of cell size. Interestingly, *P*_*Tet*_*-cdr2(1-590)* was capable of inducing Wee1 hyperphosphorylation despite a lack of clustering. *P*_*Tet*_*-cdr2(1-330)* did not induce Wee1 hyperphosphorylation, but stronger overexpression with the thiamine repressible *P3*_*nmt1*_ promoter led to Wee1 hyperphosphorylation by both Cdr2(1-330) and Cdr2(1-590) (Figure S3). For both Cdr2 truncation constructs, the catalytically inactive E177A mutant prevented hyperphosphorylation of Wee1 (Figures 5E and S3). We conclude that hyperphosphorylation of Wee1 is not sufficient to induce premature mitotic entry when Cdr2 levels increase. This result emphasizes the functional importance of Wee1 sequestration in driving early mitosis. Put together, our results show that both kinase activity and clustering are required for elevated levels of Cdr2 to induce premature mitotic entry.

**Figure 5.**
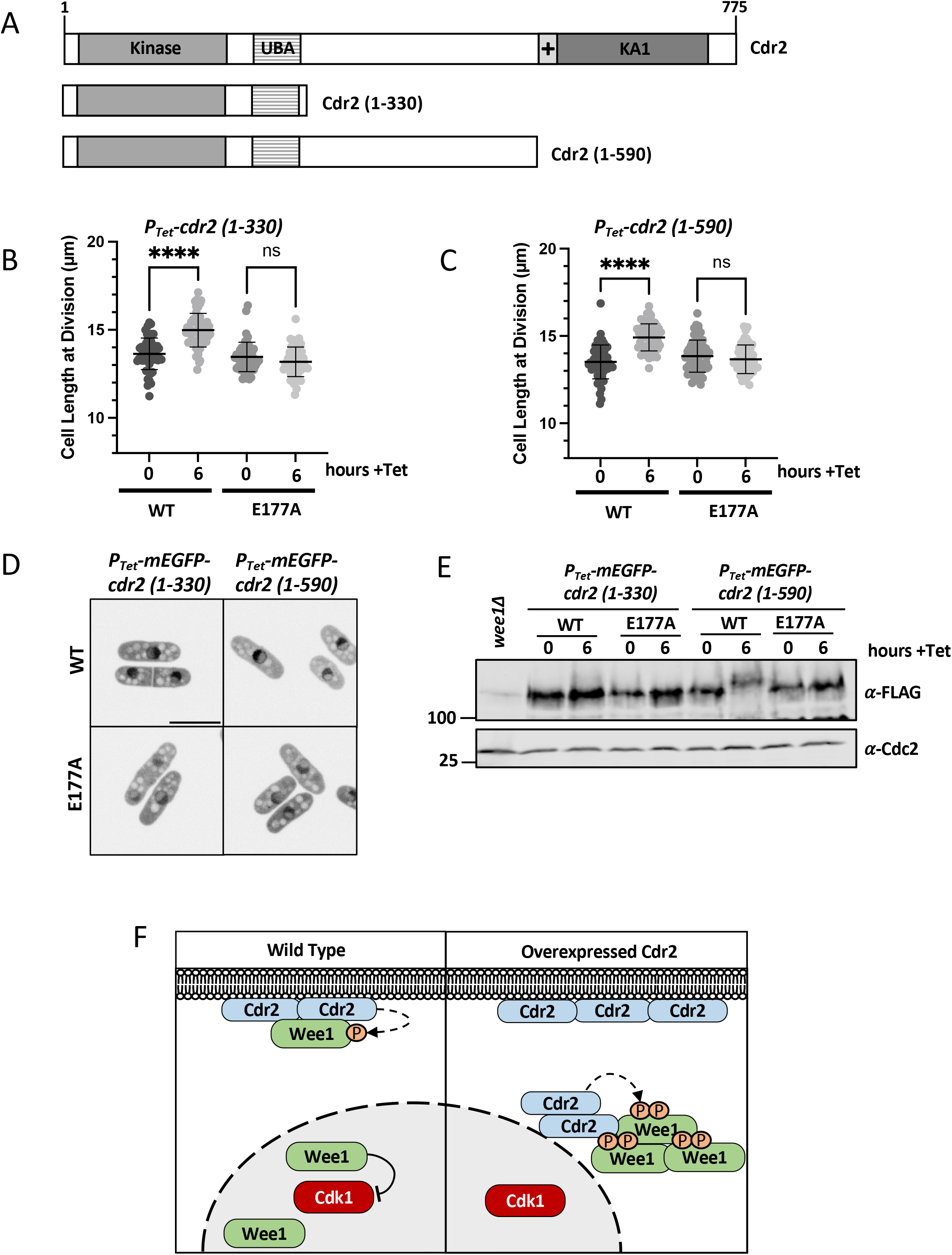
Analysis of Cdr2 truncation mutants. (A) Domain layout of Cdr2 constructs. UBA, ubiquitin-associated; KA1, Kinase associated-1; +, basic patch. (B) Cell size at division for wild type and E177A mutant versions of *P*_*Tet*_*-cdr2(1-330)*. n ≥ 50 cells. (C) Cell size at division for wild type and E177A mutant versions of *P*_*Tet*_*-cdr2(1-590)*. n ≥ 50 cells. ns, not significant (p > 0.05). ****, p < 0.0001 determined by ANOVA. (D) Localization of the indicated constructs after 6-hr _Tet_ incubation. Scale bar 10µm. (E) Western blot for Wee1-FLAG before and after induction of Cdr2 P_Tet_-GST truncations. (F) Working model for mechanism of Wee1 inhibition by over-expressed Cdr2.

In conclusion, we have shown that increased expression of Cdr2 causes early mitotic entry through a mechanism based on hyperphosphorylation and sequestration of Wee1 (Figure 5F). This phenotype means that Cdr2 overexpression versus Cdr2 loss-of-function mutations have opposite effects: _*cdr2*_*Δ* mutants delay mitotic entry leading to enlarged cells, while _*cdr2*_*+* overexpression causes premature mitotic entry leading to small cells. The mechanism that we have uncovered for upregulated Cdr2 is related but distinct from the activity of basally expressed Cdr2 at nodes. At endogenous expression levels, Cdr2 forms nodes that bring together Cdr1 and Wee1 to stimulate inhibitory phosphorylation of Wee1. In contrast, upregulated Cdr2 forms clusters away from the cell cortex and sequesters Wee1 at these sites. Concentration of Wee1 at these clusters likely prevents its interactions with Cdk1 in the nucleus. Cdr2 kinase activity is required for sequestration of Wee1, similar to the role of Cdr2 kinase activity in promoting the dwell time of Wee1 at cortical nodes (Allard *et al*., 2018). This sequestration mechanism could explain the stronger cell size defect for _*cdr2*_*Δ* cells compared to *cdr1Δ* cells (Martin and Berthelot-Grosjean, 2009; Moseley *et al*., 2009). Our results suggest that Cdr2 nodes can sequester Wee1 away from the nucleus even in the absence of Cdr1, which inhibits Wee1 catalytic activity, consistent with Wee1 localization to nodes in *cdr1Δ* cells (Allard *et al*., 2018). Thus, our work on overexpressed Cdr2 provides mechanistic insight into localization-based regulation of Wee1 and mitotic entry.

Our results also have implications for signaling pathways beyond Cdr2 and fission yeast. A growing number of protein kinases function in oligomeric clusters that contribute to their activity and regulation. Examples from multiple animal cell polarity and receptor-associated tyrosine kinase systems have revealed that clustering can promote kinase signaling (Douglass and Vale, 2005; Dickinson *et al*., 2017; Mayer and Yu, 2018). Similarly, regulated clustering of bacterial kinases such as *C. crescentus* DivJ can activate downstream signals (Saurabh *et al*., 2022). This widespread phenomenon makes kinase clustering an attractive candidate for synthetic biology approaches that seek to control signaling networks (Yoshikawa *et al*., 2021). Our work adds a new layer to this theme by showing how kinase clusters can sequester substrates to control downstream signaling, which could represent a new mechanism for these and other signaling systems.

## Materials and Methods

### Yeast Strains and Growth

Standard *Schizosaccharomyces pombe* media and methods were used (Moreno *et al*., 1991). Strains and plasmids in this study are listed in Supplemental Table S1 (listed in order that they appear in the paper). All *P*_*Tet*_ plasmids were generated using Gibson reactions (NEB HiFi Assembly Mix) from PCR products. These plasmids contain the strong *eno101* promoter with tet operons (Patterson *et al*., 2019) A linearized version of this plasmid was generated by PCR and then transformed into strains containing _Tet_R. Transformants were selected by hygromycin resistance and leucine auxotrophy. In figures 3B, 4D, 4E, and S2D, cells were cultured in YE4S (rich) media overnight, and then switched to EMM4S (minimal) media for at least 24 hours prior to induction and imaging. All other imaging was performed in YE4S.

### Overexpression of Cdr2

Strains confirmed for *P*_*Tet*_*-GST-cdr2* were grown in YE4S at 32°C for cell size measurements and Western blot sample collection, or alternatively at 25°C for live-cell fluorescent microscopy. With the exception of mEGFP-tagged constructs, all other *P*_*Tet*_*-cdr2* constructs in our study contained an N-terminal GST tag that was used to verify expression by Western blot. Cells were maintained in logarithmic phase growth at least 24 hours prior to induction. Overexpression was induced using a final concentration of 5µg/mL anhydro_Tet_racycline (_Tet_) (Sigma), from a 10mg/mL stock in DMSO (stored at -20°C). Uninduced cells in log phase were imaged for timepoint zero, and cells that were induced were seeded at OD_600_=0.05 and left to grow in the presence of Tet for 6 hours before imaging. For experiments with *wee1-50* strains, cells were treated with DMSO or _Tet_ for 2 hours at 25°C, and then were left at 25°C or shifted to 36°C for the remaining 4 hours. In Figure S3, pREP3x overexpression of Cdr2, cells containing pREP3x-6His-Cdr2 plasmid were grown in EMM4S lacking leucine and containing thiamine at 32°C. Cultures were washed into media lacking thiamine to induce expression, and then samples were collected at the indicated times.

### Western Blots

To make whole cell lysates, 2 OD_600_ units of cells were harvested at timepoint zero (prior to Tet addition) and at subsequent timepoints as indicated. Cells were pelleted, washed once with water, and pellets were flash frozen in liquid nitrogen. Samples were lysed by bead beating with Mini-beadbeater-16 in SDS-PAGE sample buffer including protease and phosphatase inhibitors (15% glycerol, 4.5% SDS, 97.5mM Tris pH6.8, 10% 2-mercaptoethanol, 50mM ?-glycerophosphate, 50mM sodium fluoride, 5mM sodium orthovanadate, 1x EDTA-free protease inhibitor cocktail (Sigma Aldrich)), and then incubated at 99°C for 5 minutes. Following brief centrifugation, clarified lysate was separated by SDS-PAGE and transferred to nitrocellulose using Trans-blot Turbo Transfer System (Bio-Rad). Proteins were probed with antibodies against FLAG M2 (Sigma), Cdc2 (Santa Cruz Biotechnologies SC-53217), and Wee1 (Allard *et al*., 2018). Blots were imaged on a LiCor Odyssey CLx. For quantification in Figure 1C, three independently grown biological replicate cultures were prepared for each condition and analyzed in the Western blot as shown. Band intensities were measured with ImageStudioLite software and normalized to levels of Cdr2-9gly-5xFLAG expressed from the endogenous promoter and chromosomal locus. The *P*_*Tet*_*-cdr2-9Gly-5xFLAG* strain in this experiment was a second copy expressed in addition to with the untagged, endogenous Cdr2. Data were graphed using Prism9 Graphpad.

### Lambda Phosphatase

The protocol was adapted from previous work (Lucena *et al*., 2017) using phosphatase buffer prepared to 1x and containing 1 mM MnCl_2_. In brief: 2 OD_600_ units of cells were pelleted, flash frozen in nitrogen, lysed by bead beating in 200µL phosphatase buffer with glass beads for 1 minute, placed on ice for 1 minute, and then centrifuged at 15,000*g* for 1 minute. For each reaction, 20µL of this lysate was mixed with 800U lambda phosphatase (New England Biolabs) or untreated. These reactions were incubated for 30 minutes at 30° C. Reactions were stopped by addition of 2x SDS-PAGE sample buffer (30% glycerol, 9% SDS, 195mM Tris pH 6.8, 15% 2-mercaptoethanol) and boiled for 5 minutes before analyzing by SDS-PAGE as described above.

### Widefield Microscopy

Images for cell length measurements (Figure 2, 5B-C) were captured at room temperature on a DeltaVision imaging system comprised of an Olympus IX-71 inverted wide-field microscope, a Photometrics Cool-SNAP HQ2 camera, an Insight solid-state illumination unit, and a 1.42 NA Plan Apo 60x oil objective. Cells were grown as specified above and imaged after addition of Blankophor cell wall stain to identify dividing, septated cells. 1,6-hexanediol-treated cells were also imaged on this system (Figure 4C, S2C). These cells were grown and induced with Tet in YE4S as described, washed for 20 minutes into fresh YE4S, then incubated another 10 minutes in the presence or absence of 5% (w/v) 1,6-hexanediol dissolved in media.

### Spinning-Disc Confocal Microscopy

Fluorescent live-cell microscopy (Figures 3, 4A-B, 4D-E, 5D, S2A-B, and S2D) was performed at room temperature on a Yokogawa CSU-WI imaging system, which was equipped with a Nikon Eclipse Ti2 inverted microscope, a 100× 1.45 NA CFI Plan Apochromat Lambda objective lens (Nikon); 405-, 488-, and 561-nm laser lines; and a photometrics Prime BSI camera. Due to fluorescence of anhydro_Tet_racycline, imaging of fluorescently tagged proteins was performed on cells that had been washed into fresh _Tet_-free media for 20 minutes prior to imaging.

### Cell Length Measurements and Statistics

Cell length measurements were made using Line Tool in FIJI Image J (Schindelin *et al*., 2012) on images of cells stained with the cell wall dye Blankophor. The resulting data were graphed and statistically analyzed in Prism9 Graphpad. Cell length measurements in Figure 2E were compared by Welch’s t test. All other cell length measurements were compared by one-way ANOVA followed by Tukey’s multiple comparison test, which compares each mean within an experiment to each other.

## Acknowledgements

We thank members of the Moseley laboratory for helpful discussions and comments on the manuscript. We thank the Nurse laboratory for sharing *P*_*Tet*_ strains and plasmids, Scott Curran for experimental advice, and the Biomolecular Targeting Core (bioMT) for equipment (P20-GM113132). This work was supported by grants from the National Institute of General Medical Sciences (R01GM099774 and R01GM133856) to J.B.M.

## Figure Legends

Figure S1. *P*_*Tet*_*-cdr2* induces hyperphosphorylation of untagged Wee1. (A) Induction of *P*_*Tet*_*-* _*cdr2*_*(T166A)* does not lead to Wee1-FLAG hyperphosphorylation. (B) Whole-cell extracts were separated by SDS-PAGE and Western blotted against endogenous Wee1. Asterisk marks background band. (C) Western blot as in panel B, but in _*cdr2*_*Δ cdr1Δ* cells.

Figure S2. Localization of wild type and E177A Cdr2 constructs. (A) P_Tet_-mEGFP-Cdr2(E177A) colocalized with Cdr2(E177A)-mCherry. (B) Wee1-mNG and Cdr2(E177A)-mCherry in *P*_*Tet*_*-* _*cdr2*_*(E177A)* induced conditions. (C) Treatment of endogenously expressed *mEGFP-*_*cdr2*_ cells with 5% 1,6-hexanediol disrupts cortical nodes. Middle focal plane. (D) Localization of Wee1-mNG and Cut11-mCherry after P_Tet_-GST-Cdr2 induction. All scale bars are 10 µm. Single channel images are shown with inverted LUT. Panels A, B, and D are maximum intensity projections of 0.5 µm-spaced focal planes covering middle 1 µm of cells.

Figure S3. Overexpression of Cdr2 truncations by the strong *P3*_*nmt1*_ promoter. (A) Western blot for Wee1-FLAG after induction of pREP3x-Cdr2(1-330) and pREP3x-cdr2(1-330; E177A). (B) Western blot of Wee1-FLAG after induction of pREP3x-Cdr2(1-590) and pREP3x-cdr2(1-590; E177A).

## Notes

### Competing Interest Statement

The authors have declared no competing interest.

