## Supplemental Figures 1-3 and Table 1 for "Regulation of cell size and Wee1 by elevated levels of Cdr2"

Figure S1

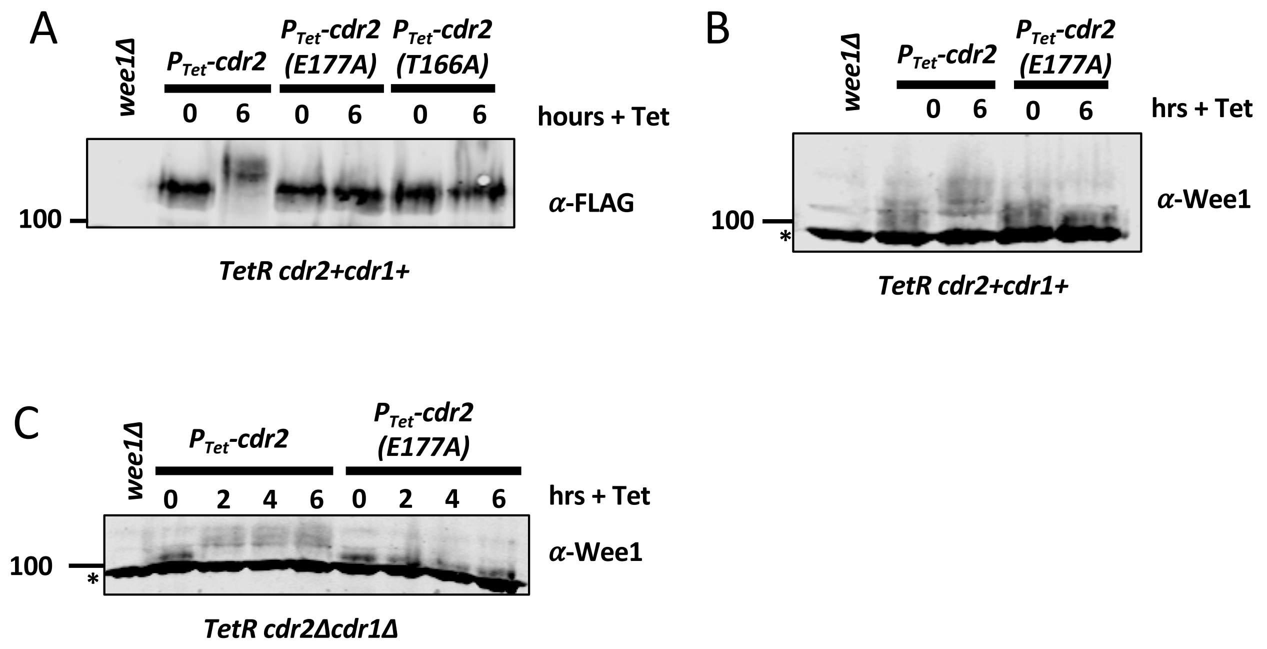
A
wee1Δ
P<sub>Tet</sub>-cdr2
P<sub>Tet</sub>-cdr2 (E177A)
P<sub>Tet</sub>-cdr2 (T166A)
0 6 0 6 0 6
hours + Tet
α-FLAG
100
TetR cdr2+cdr1+
B
wee1Δ
P<sub>Tet</sub>-cdr2
P<sub>Tet</sub>-cdr2 (E177A)
0 6 0 6
hrs + Tet
α-Wee1
100
*
TetR cdr2+cdr1+
C
wee1Δ
P<sub>Tet</sub>-cdr2
P<sub>Tet</sub>-cdr2 (E177A)
0 2 4 6 0 2 4 6
hrs + Tet
α-Wee1
100
*
TetR cdr2Δcdr1Δ

Figure S2

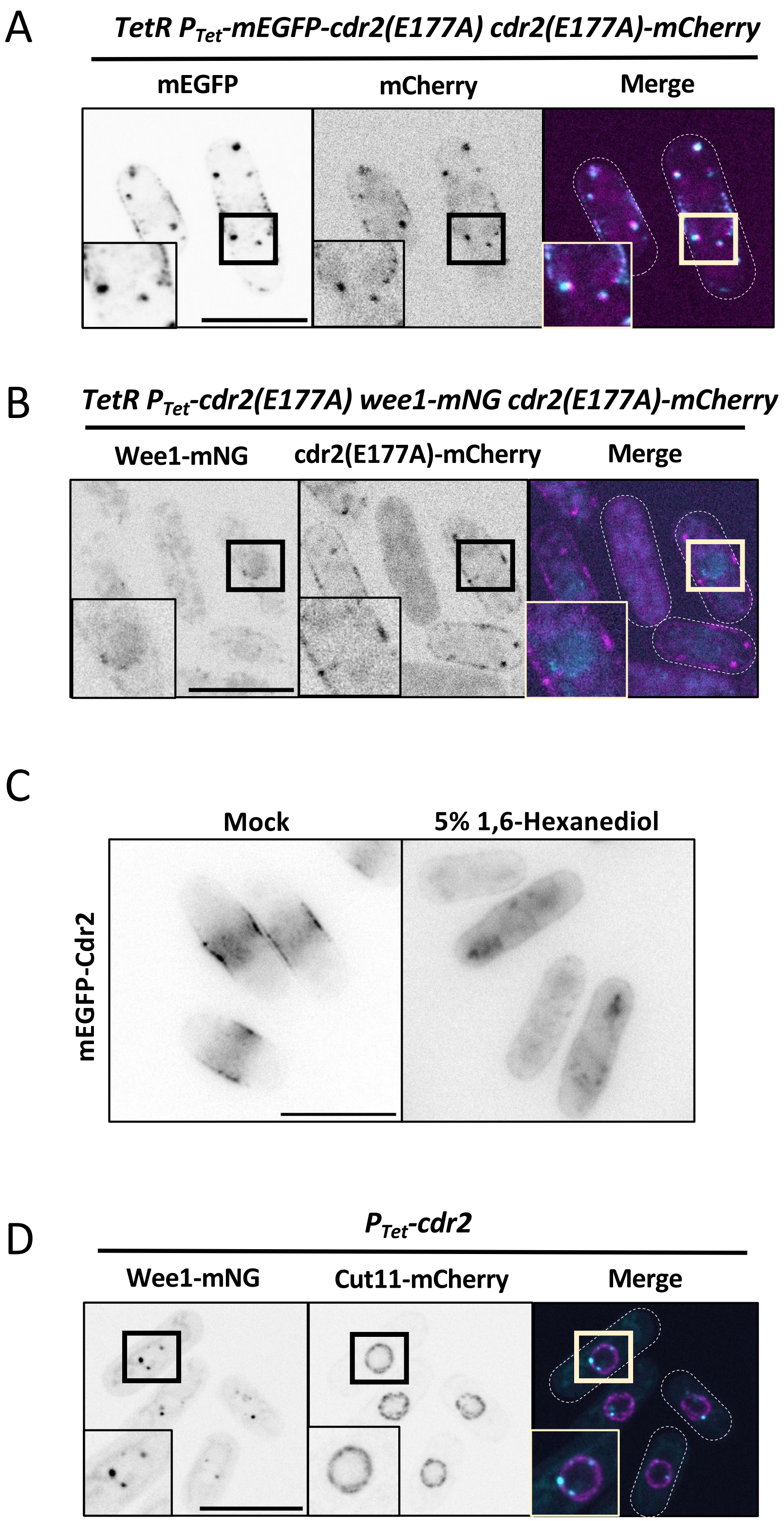
A
TetR PTet-mEGFP-cdr2(E177A) cdr2(E177A)-mCherry
mEGFP
mCherry
Merge
B
TetR PTet-cdr2(E177A) wee1-mNG cdr2(E177A)-mCherry
Wee1-mNG
cdr2(E177A)-mCherry
Merge
C
Mock
5% 1,6-Hexanediol
mEGFP-Cdr2
D
PTet-cdr2
Wee1-mNG
Cut11-mCherry
Merge

Figure S3

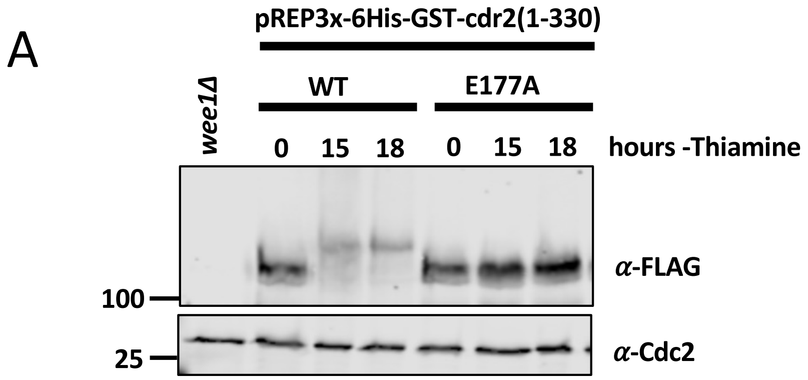
A
pREP3x-6His-GST-cdr2(1-330)
wee1Δ
WT
E177A
0
15
18
0
15
18
hours -Thiamine
α-FLAG
100
α-Cdc2
25

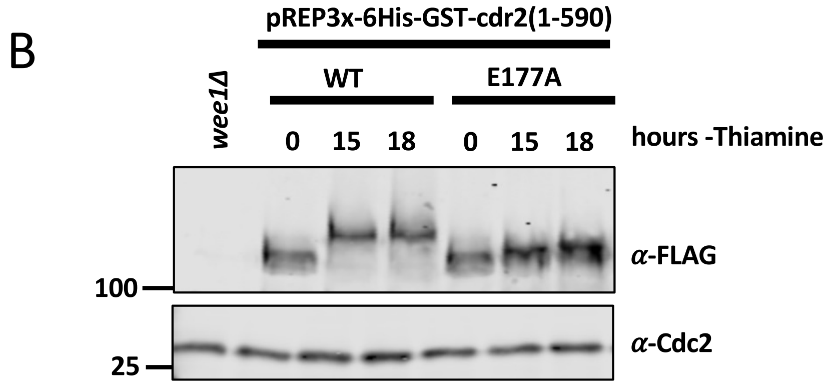
B
pREP3x-6His-GST-cdr2(1-590)
wee1Δ
WT
E177A
0
15
18
0
15
18
hours -Thiamine
α-FLAG
100
α-Cdc2
25

### Table S1

| Strain Name | Genotype | Source |
|---|---|---|
| JM7124 | *h- TetR-ura4+* | Paul Nurse Lab (JP521) |
| JM7525 | *wee1-9gly5Flag::hphR ura4-D18::TetR-ura4+ leu1::enoTet-GST-cdr2-hphR* | This Study |
| JM7526 | *wee1-9gly5Flag::hphR ura4-D18::TetR-ura4+ leu1::enoTet-GST-cdr2(E177A)-hphR* | This Study |
| JM777 | *wee1Δ::ura4+ ura4-D18 leu1-32 h-* | Lab Collection |
| JM6129 | *cdr2-9gly-5FLAG::kanR h+* | Lab Collection |
| JM7481 | *ura4-D18::TetR-ura4+ leu1::enoTet-cdr2-9gly5Flag* | This Study |
| JM7585 | *cdr2Δ::natR nim1Δ::kanMX6 ura4-D18::TetR-ura4+ leu1::enoTet-GST-cdr2-hphR wee1-9gly5Flag::hphR* | This Study |
| JM7595 | *cdr2Δ::natR nim1Δ::kanMX6 ura4-D18::TetR-ura4+ leu1::enoTet-GST-cdr2(E177A)-hphR wee1-9gly5Flag::hphR* | This Study |
| JM7473 | *ura4-D18::TetR-ura4+ leu1::enoTet-GSTcdr2-hphR h+* | This Study |
| JM7474 | *ura4-D18::TetR-ura4+ leu1::enoTet-GSTcdr2(E177A)-hphR h+* | This Study |
| JM7402 | *cdr2Δ::natR cdr1Δ::kanMX6 ura4-D18::TetR-ura4+ leu1::enoTet-GST-cdr2-hphR* | This Study |
| JM7403 | *cdr2Δ::natR cdr1Δ::kanMX6 ura4-D18::TetR-ura4+ leu1::enoTet-GST-cdr2(E177A)-hphR* | This Study |
| JM7675 | *wee1-50 ura4-D18::TetR-ura4+ leu1::enoTetGST-cdr2-hphR* | This Study |
| JM7680 | *ura4-D18::TetR-ura4+ leu1::enoTet-mEGFP-cdr2-hphR h+* | This Study |
| JM7691 | *ura4-D18::TetR-ura4+ leu1::enoTet-mEGFP-cdr2(E177A)-hphR h+* | This Study |
| JM7574 | *ura4-D18::TetR-ura4+ leu1::enoTet-GST-cdr2-hphR wee1-yomNeonGreen::hphR* | This Study |
| JM7575 | *ura4-D18::TetR-ura4+ leu1::enoTet-GST-cdr2(E177A)-hphR wee1-yomNeonGreen::hphR* | This Study |
| JM7712 | *ura4-D18::TetR-ura4+ leu1::enoTet-mEGFP-cdr2-hphR cdr2-mCherry::natR* | This Study |
| JM7583 | *cdr2-mCherry::natR wee1-yomNeonGreen::hphR ura4-D18::TetR-ura4+ leu1::enoTet-GST-cdr2-hphR* | This Study |
| JM7709 | *cdr1-gammaGFP::hphR ura4-D18::TetR-ura4+ leu1::enoTet-GST-cdr2-hphR* | This Study |
| JM7710 | *cdr1-gammaGFP::hphR ura4-D18::TetR-ura4+ leu1::enoTet-GST-cdr2(E177A)-hphR* | This Study |
| JM7671 | *arf6-yomNeonGreen::hphR ura4-D18::TetR-ura4+ leu1::enoTet-GST-cdr2-hphR* | This Study |
| JM7672 | *arf6-yomNeonGreen::hphR ura4-D18::TetR-ura4+ leu1::enoTet-GST-cdr2(E177A)-hphR* | This Study |
| JM7655 | *wee1-yomNeonGreen::hphR ura4-D18::TetR-ura4+ leu1::enoTet-GST-cdr2-hphR sur4-mCherry::natR* | This Study |
| JM7662 | *wee1-yomNeonGreen::hphR ura4-D18::TetR-ura4+ leu1::enoTet-GST-cdr2-hphR sad1-mCherry::natR* | This Study |
| JM7553 | *ura4-D18::TetR-ura4+ leu1::enoTet-GST-cdr2(1-330aa)-hphR* | This Study |
| JM7554 | *ura4-D18::TetR-ura4+ leu1::enoTet-GST-cdr2(1-330aa; E177A)-hphR* | This Study |
| JM7581 | *ura4-D18::TetR-ura4+ leu1::enoTet-GST-cdr2(1-590aa)-hphR* | This Study |
| JM7582 | *ura4-D18::TetR-ura4+ leu1::enoTet-GST-cdr2(1-590aa; E177A)-hphR* | This Study |
| JM7681 | *ura4-D18::TetR-ura4+ leu1::enoTet-mEGFP-cdr2(1-330aa)-hphR h+* | This Study |
| JM7682 | *ura4-D18::TetR-ura4+ leu1::enoTet-mEGFP-cdr2(1-590aa)-hphR h+* | This Study |
| JM7692 | *ura4-D18::TetR-ura4+ leu1::enoTet-mEGFP-cdr2(1-330aa; E177A)-hphR h+* | This Study |
| JM7693 | *ura4-D18::TetR-ura4+ leu1::enoTet-mEGFP-cdr2(1-590aa; E177A)-hphR h+* | This Study |
| JM7563 | *ura4-D18::TetR-ura4+ leu1::enoTet-GST-cdr2(1-330aa)-hphR wee1-9gly5Flag::natR* | This Study |
| JM7566 | *ura4-D18::TetR-ura4+ leu1::enoTet-GST-cdr2(1-330aa; E177A)-hphR wee1-9gly5Flag::natR* | This Study |
| JM7665 | *ura4-D18::TetR-ura4+ leu1::enoTet-GST-cdr2(1-590aa)-hphR wee1-9gly5FLAG::hphR* | This Study |
| JM7624 | *ura4-D18::TetR-ura4+ leu1::enoTet-GST-cdr2(1-590aa; E177A)-hphR wee1-9gly5Flag::hphR* | This Study |
| JM7576 | *ura4-D18::TetR-ura4+ leu1::enoTet-GST-cdr2(T166A)-hphR wee1-9gly5Flag::hphR* | This Study |
| JM7713 | *ura4-D18::TetR-ura4+ leu1::enoTet-mEGFP-cdr2(E177A)-hphR cdr2(E177A)-mCherry::natR* | This Study |
| JM7729 | *ura4-D18::TetR-ura4+ leu1::enoTet-GST-cdr2(E177A)-hphR wee1-yomNeonGreen::hphR cdr2(E177A)-mCherry::natR* | This Study |
| JM2576 | *mEGFP-cdr2* | Lab Collection |
| JM7654 | *wee1-yomNeonGreen::hphR ura4-D18::TetR-ura4+ leu1::enoTet-GST-cdr2-hphR cut11-mCherry::kanMX6* | This Study |
| JM7562 | *cdr2Δ::natR nim1Δ::kanMX6 wee1-9gly5FLAG::hphR ura4-D18 leu1-32* | This Study |

| Plasmid Name | Description | Source |
|---|---|---|
| pJM210 | pREP3X | |
| pJM1574 | pREP3x-6HisGST-cdr2(1-330) | |
| pJM1575 | pREP3x-6HisGST-cdr2(1-590) | |
| pJM1593 | JPp135 | Paul Nurse Lab |
| pJM1595 | pREP3x-6HisGST-cdr2 330(E177A) | |
| pJM1622 | pREP3x-6hisGST-cdr2(590aa)(E177A) | |
| pJM1607 | eno101P-3xTetR-GST-Cdr2-Tadh1 | |
| pJM1608 | eno101P-3xTetR-GST-cdr2(E177A)-Tadh1 | |
| pJM1609 | eno101P-3xTetR-GST-cdr2(T166A)-Tadh1 | |
| pJM1610 | eno101P-3xTetR-GST-Cdr2(aa1-330)-Tadh1 | |
| pJM1611 | eno101P-3xTetR-GST-cdr2(E177A) (aa1-330)-Tadh1 | |
| pJM1620 | eno101P-3xTetR-GSTcdr2(590aa)-Tadh1 | |
| pJM1621 | eno101P-3xTetR-GSTcdr2(590aa)(E177A)-Tadh1 | |
| pJM1624 | eno101P-3xTetR-Cdr2-9gly5FLAG-Tadh1 | |
| pJM1631 | eno101P-3xTetR-mEGFP-Cdr2-Tadh1 | |
| pJM1632 | eno101P-3xTetR-mEGFP-cdr2(330 aa)-Tadh1 | |
| pJM1633 | eno101P-3xTetR-mEGFP-cdr2(590 aa)-Tadh1 | |
| pJM1634 | eno101P-3xTetR-mEGFP-cdr2(E177A)-Tadh1 | |
| pJM1635 | eno101P-3xTetR-mEGFP-cdr2(E177A)(330 aa)-Tadh1 | |
| pJM1636 | eno101P-3xTetR-mEGFP-cdr2(E177A)(590 aa)-Tadh1 | |
